## Supplementary Information for "Hyphal compartmentalization and sporulation in *Streptomyces* require the conserved cell division protein SepX"

**Affiliations**
[1] Department of Molecular Microbiology, John Innes Centre, Norwich Research Park, Norwich NR4-7HU UK
[2] Department of Cell and Developmental Biology, John Innes Centre, Norwich Research Park, Norwich, NR4-7HU, UK
[3] Département de Microbiologie, Infectiologie et Immunologie, Université de Montréal, Pavillon Roger-Gaudry, 2900, boulevard Édouard-Montpetit, C.P. 6128, Succursale Centre-ville, Montréal, QC H3C 3J7, Canada

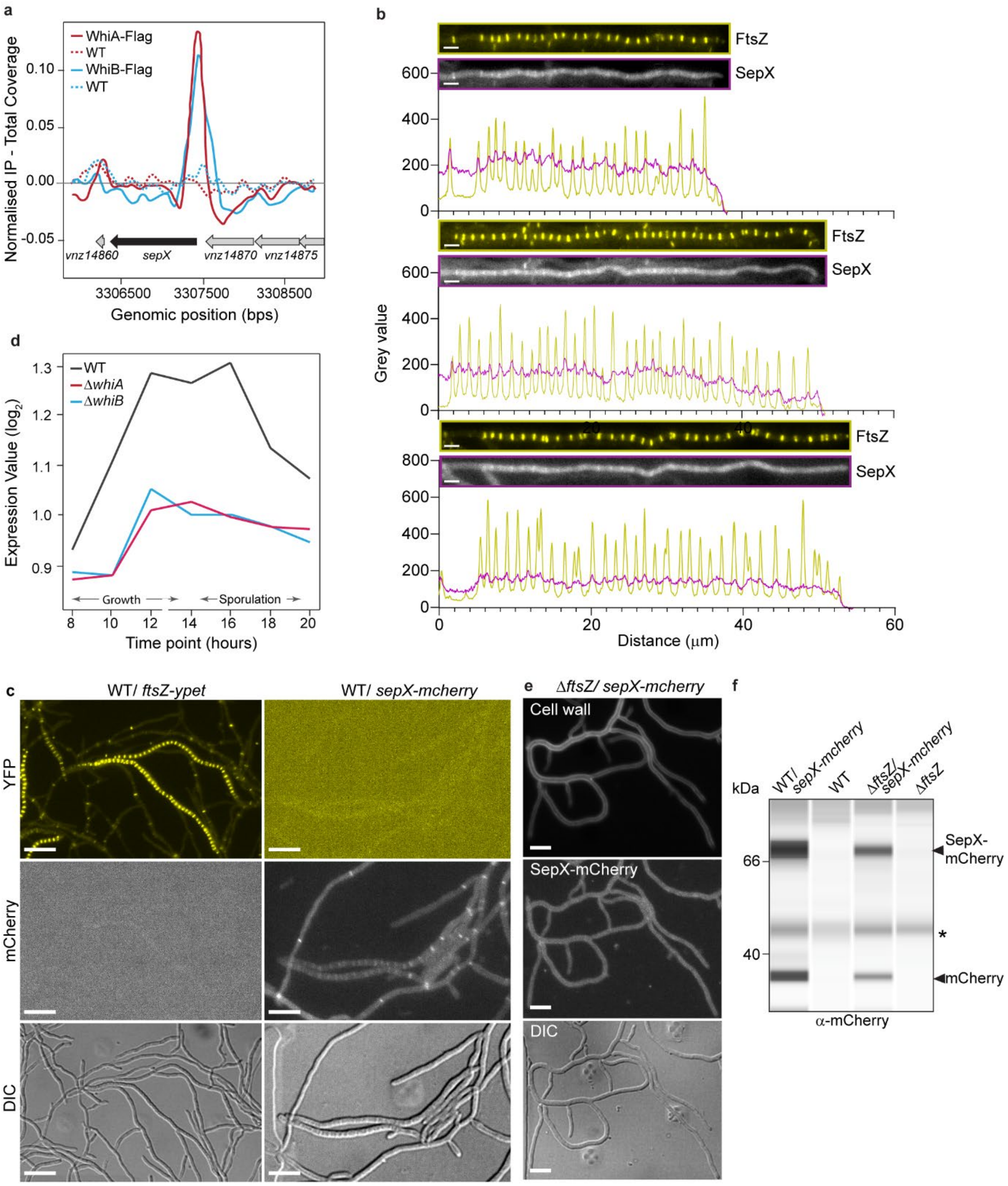

**Supplementary Figure 1: SepX is developmentally regulated and requires FtsZ for localization. (a)** *sepX* (*vnz_14865*) is a WhiAB-target as identified by ChIP-seq. Enrichment (normalized IP – total coverage) in strains expressing either WhiA-FLAG (solid red line) or WhiB-FLAG (solid blue line) but not in the untagged wildtype controls (dashed red and dashed blue lines) was observed upstream of the *sepX* gene *in vivo*[14,15]. **(b)** Additional fluorescence intensity profiles of FtsZ-YPet and SepX-mCherry in sporulating hyphae shown in Supplementary Movie 1. Scale bars: 2 μm. Data shown are representatives of four independent hyphae. Source data are provided as a Source Data file. **(c)** Control images showing the absence of spectral bleedthrough and the specificity of the FtsZ-YPet and SepX-mCherry fluorescence signal using the same illumination settings (strains LUV015, MB1124). Scale bars: 5 μm. Data shown are representatives of at least 10 images. **(d)** WhiA and WhiB co-activate the transcription of *sepX.* Data represent transcriptomic data during submerged sporulation in wild-type *S. venezuelae* (black line); the congenic *whiA* mutant (red line); and the congenic *whiB* mutant (blue line)[14,15,19]. The x-axis indicates the age of the culture in hours, and the y-axis indicates the per-gene normalized transcript abundance (Expression Value $\log_2$). **(e)** Microscopic analysis of SepX-mCherry distribution in the $\Delta$*ftsZ* mutant background (MB1082). Hyphal cell wall was visualized using 0.25 mM HADA. DIC, differential interference contrast. Scale bar: 5 $\mu$m. Data shown are representatives of at least 10 images. **(f)** Virtual Western blot showing SepX-mCherry abundance in the wildtype (MB1124) and the $\Delta$*ftsZ* mutant (MB1082) compared to the corresponding untagged strains. Automated western blot analysis was performed in biological duplicate. Equal amounts of protein lysate were loaded and SepX-mCherry was detected using $\alpha$-mCherry antibody. Asterisk denotes an unspecific signal detected in all samples.

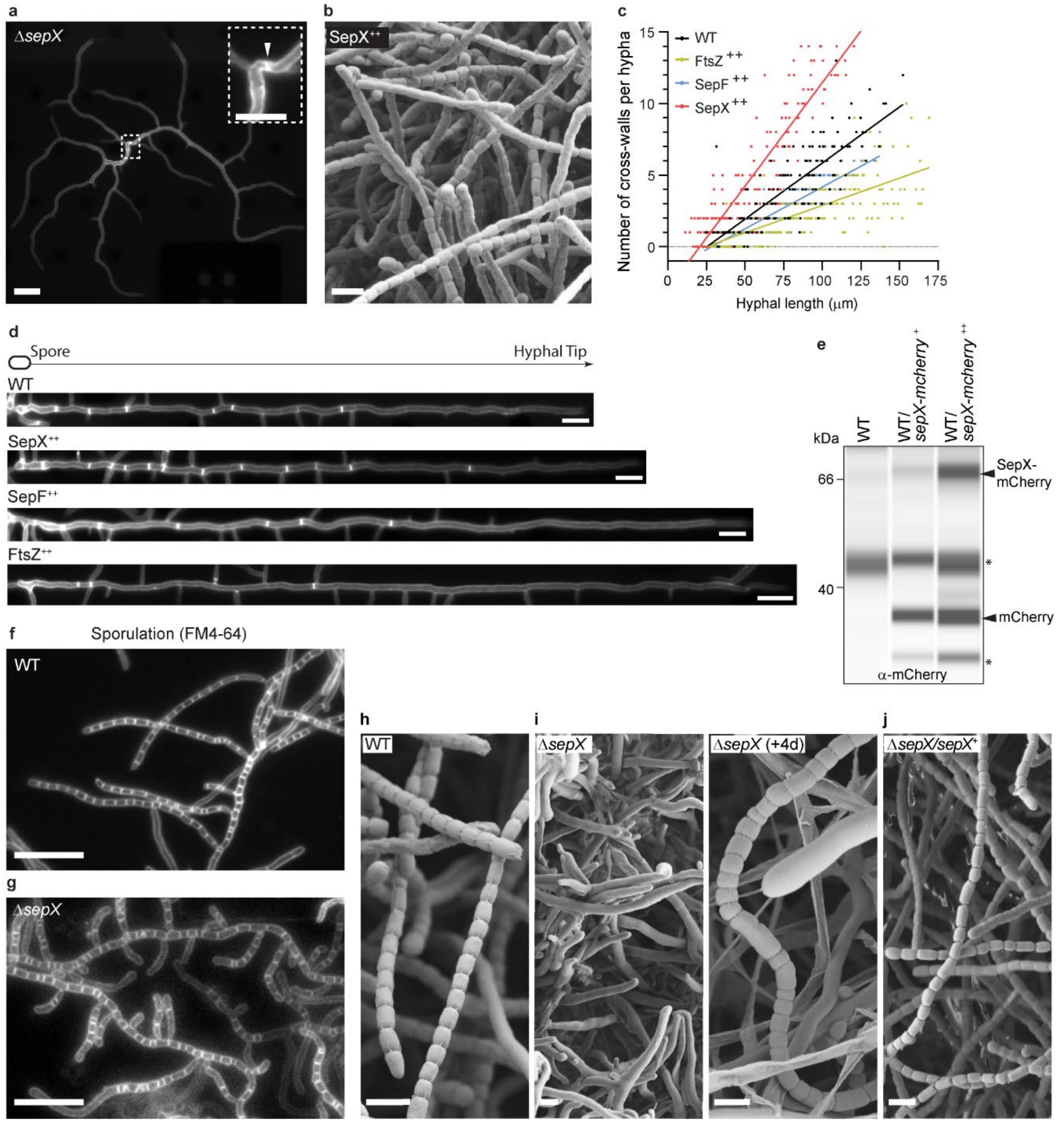

**Supplementary Figure 2: SepX is a determinant of cross-wall formation and is required for regular sporulation. (a)** Representative image of HADA-stained hyphae emerging from a Δ*sepX* spore (small dashed box) Scale bar: 10 μm. Occasionally, a division septum can be detected close to the mother spore (magnified region in inset). Scale bar: 5 μm. Experiments were performed in triplicate. **(b)** Cryo-SEM image of spore chains produced by wild-type *S. venezuelae* constitutively expressing *sepX* (MB168, SepX$^{++}$). Scale bar: 2 μm. (Representative of least 7 images) **(c)** Quantification of cross-wall frequency in strains constitutively expressing *sepX* (SepX$^{++}$, MB168), *ftsZ* (FtsZ$^{++}$, MB127) or *sepF* (SepF$^{++}$, SS414) compared to the wildtype (WT). Strains were allowed to germinate and grow in the presence of 0.25 mM HADA to visualize cross-walls. Quantification is based on biological triplicate experiments per strain. Hyphae emerging from at least 20 spores per replicate were analyzed. Solid line represents simple linear regression. **(d)** Representative images of straightened hyphae showing the distribution of cross-walls in the same strains analyzed in (c). Scale bars: 5 μm. Experiments were performed in triplicates. **(e)** Virtual Western blot showing SepX-mCherry levels when produced from the native promoter (+) or a constitutive promoter (++). Equal amounts of proteins were loaded and SepX-mCherry was detected using an $\alpha$-mCherry antibody. Asterisks denote unspecific signals and cleavage products. Shown are representative data from duplicate experiments. **(f)** and **(g)** Fluorescent micrographs of sporulating wild-type (WT) and Δ*sepX* (SV55) hyphae stained with FM4-64 to visualize sporulation septa. Scale bar: 10 $\mu$m. Shown are representative hyphae of at least 5 images. **(h-j)** Cryo-SEM images showing sporulation septation in hyphae of the wildtype (WT, h), the Δ*sepX* mutant (SV55, i) and the complemented mutant (MB181, j). Strains were grown on solid MYM and colonies imaged after three days (representative of at least seven images). The Δ*sepX* mutant does not sporulate after three days but eventually forms irregular spore chains after additional four days of incubation. Scale bars: 2 $\mu$m.

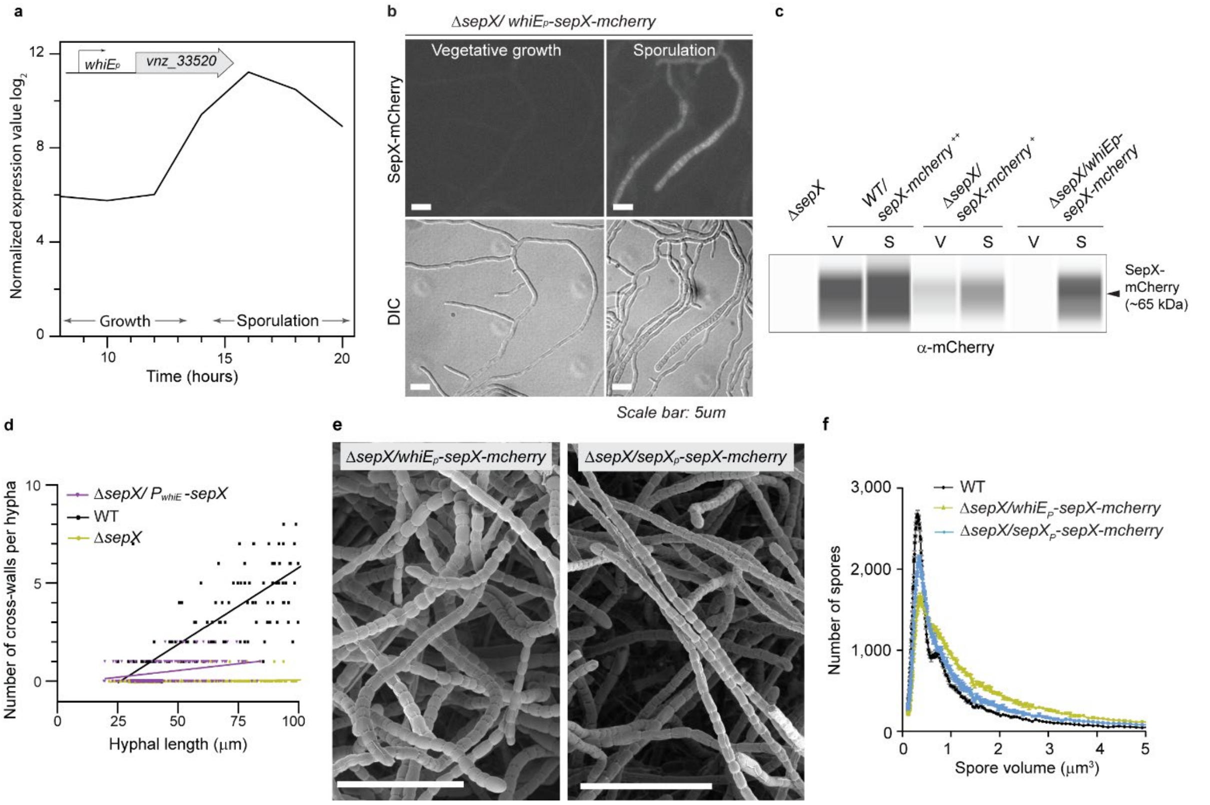

**Supplementary Figure 3: Sporulation-specific expression of *sepX* results in a largely absence of cross-walls but normal sporulation septation. (a)** Expression profile showing the sporulation-specific activity of *whiE*$_P$-driven gene expression over the *S. venezuelae* life cycle based on microarray data for *vnz_33520*. Transcriptomic source data was obtained from Al-Bassam et al. (2014)[1]. **(b)** Microscopic analysis of SepX-mCherry localization following expression from the *whiE* promoter in the Δ*sepX* mutant background (MB1120). Septal localization of SepX-mCherry is only observed during sporulation. Scale bars: 5 μm. Shown are representative of at least five images. Source data are provided as a Source Data file. **(c)** Virtual Western blot showing the abundance of SepX-mCherry during vegetative growth (V) and sporulation (S). SepX-mCherry levels were determined in the wildtype expressing *sepX* from the *ermE** constitutive promoter (++, MB1124) and in the Δ*sepX* mutant (SV55), either from the native promoter (+, MB171) or the *whiE*-promoter (MB1120). SepX-mCherry was first immunoprecipitated from whole cell lysates. Equal total protein from each strain was used as input for the immunoprecipitation and equal volume subsequently loaded for Western analysis following enrichment. Protein levels were analyzed using an α-mCherry antibody. Experiments were performed in biological duplicate. **(d)** Quantification of cross-wall frequency in the Δ*sepX* mutant expressing *sepX* from the *whiE*-promoter (MB1120). Experiments were performed as described above. Hyphae of at least 20 spores per biological replicate (n=3) were analyzed. For comparison, cross-wall distribution of the wildtype (WT) and the Δ*sepX* mutant (SV55) from Figure 2c are shown again. Solid line represents simple linear regression. **(e)** Cryo-SEM images of spore chains in the Δ*sepX* mutant complemented with *sepX-mcherry* expressed from the *whiE* promoter (left, MB1120) or the native promoter (right, MB171). Scale bar: 10 μm. Shown are representative of at least seven images per strain. **(f)** Spore volume frequency distributions (normalized to total count, n=100,000) of the wildtype (WT) and the Δ*sepX* mutant complemented with *sepX-mCherry* expressed from the *whiE* promoter (MB1120) or its native promoter (MB171). Shown is the mean ± s.e.m. of three biological replicates per strain. Source data are provided as a Source Data file.

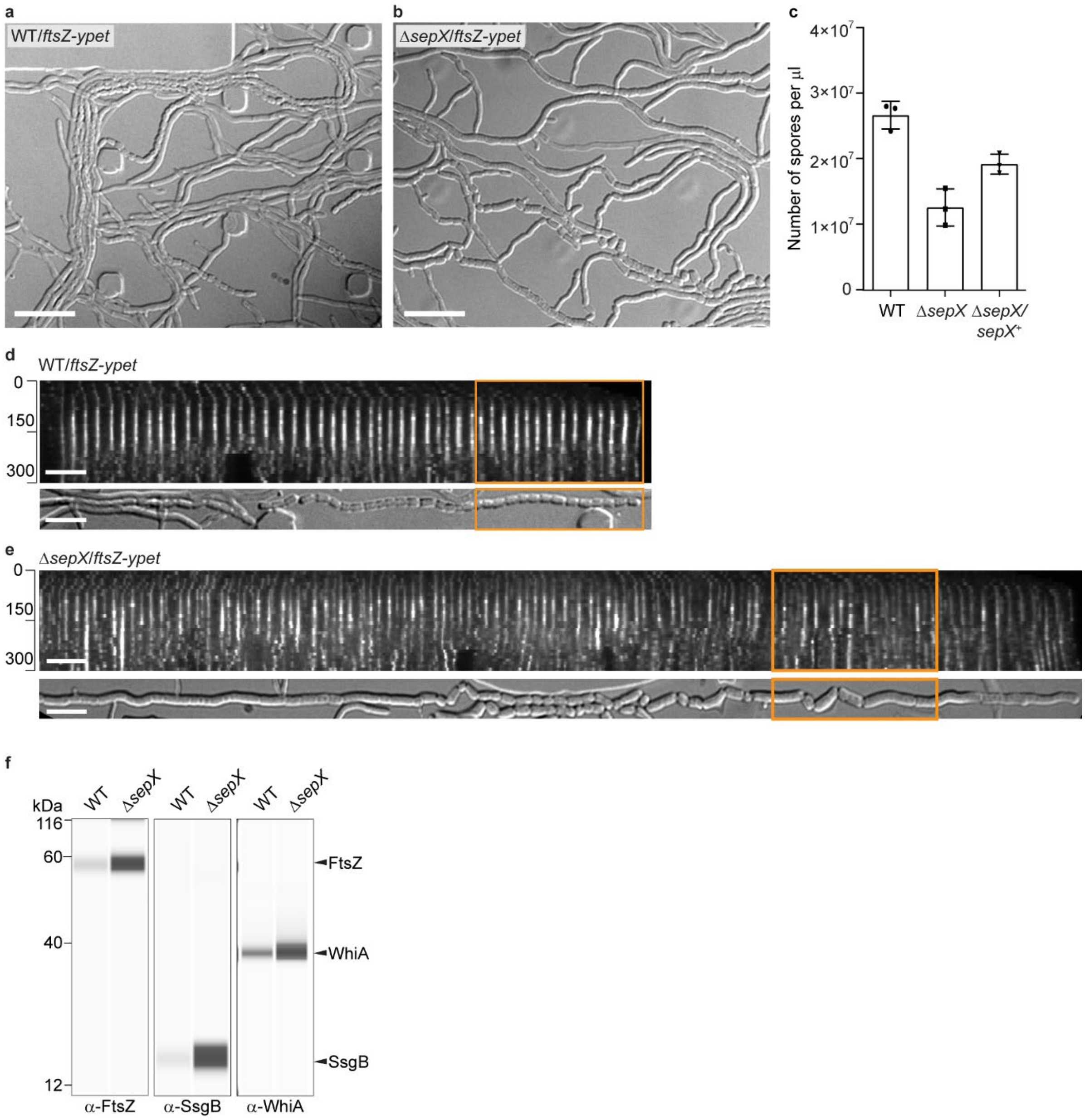

**Supplementary Figure 4: SepX is required for wildtype-like sporulation. (a)** and **(b)** are differential interference contrast (DIC) images corresponding to the fluorescent micrographs shown in Figure 4a and b. Note that the images show the outcome of the sporulation septation process in (a) the wildtype (SS12) and (b) the Δ*sepX* mutant (MB180). Scale bars: 10 µm. **(c)** Number of spores produced by the wildtype (WT), the Δ*sepX* mutant (SV55) and the complemented mutant (MB181). Shown is the mean ± s.e.m. Source data are provided as a Source Data file. **(d)** and **(e)** Representative kymographs of FtsZ-YPet fluorescence and the corresponding DIC images obtained from the fluorescence time-lapse image series (n=5 per strain) of wild-type (SS12) and *sepX*-deficient hyphae (MB180). The orange boxes indicate the subsections shown in Figure 4c and 4d. Scale bars: 5 µm. **(f)** Virtual Western blot showing the abundance of FtsZ, SsgB and WhiA in the wildtype (SS12) and the Δ*sepX* mutant (MB180) during sporulation. Samples were taken from sporulating cultures and proteins were detected using α-FtsZ, α-SsgB and α-WhiA polyclonal antibodies respectively. The data shown here are representative for experiments performed in duplicate or in triplicate (c).

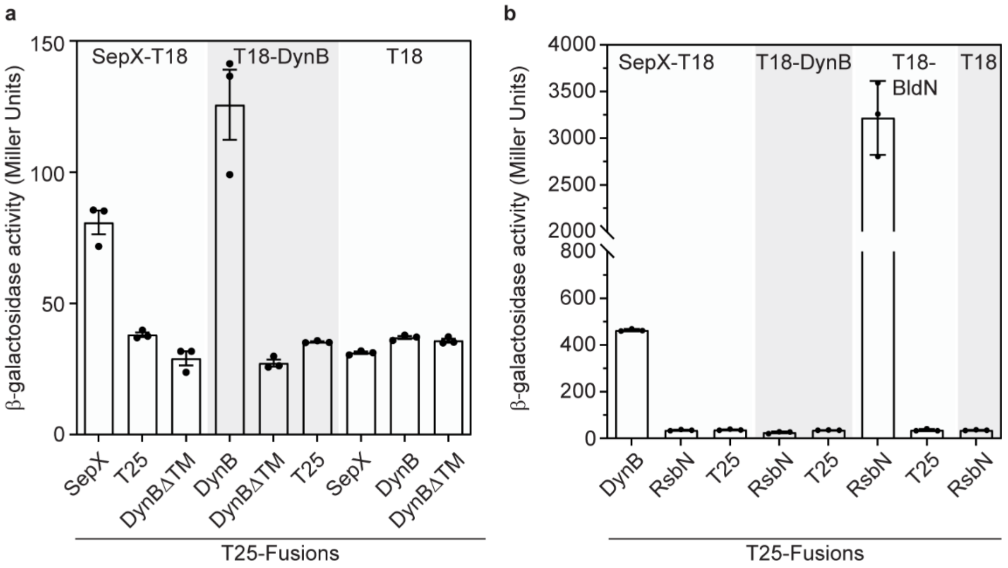

**Supplementary Figure 5: SepX specifically interacts with DynB via the DynB transmembrane segment.** Additional bacterial two-hybrid interaction studies. β-galactosidase activity was obtained from three biological replicates showing in **(a)** the self-interaction of SepX and loss of interaction with DynB carrying a deletion of the two transmembrane segments (DynBΔTM) and in **(b)** that SepX specifically interacts with DynB but not with other membrane proteins like BldN. Data points in (a) and (b) present mean values +/- s.e.m. Each interaction was assayed in triplicate.

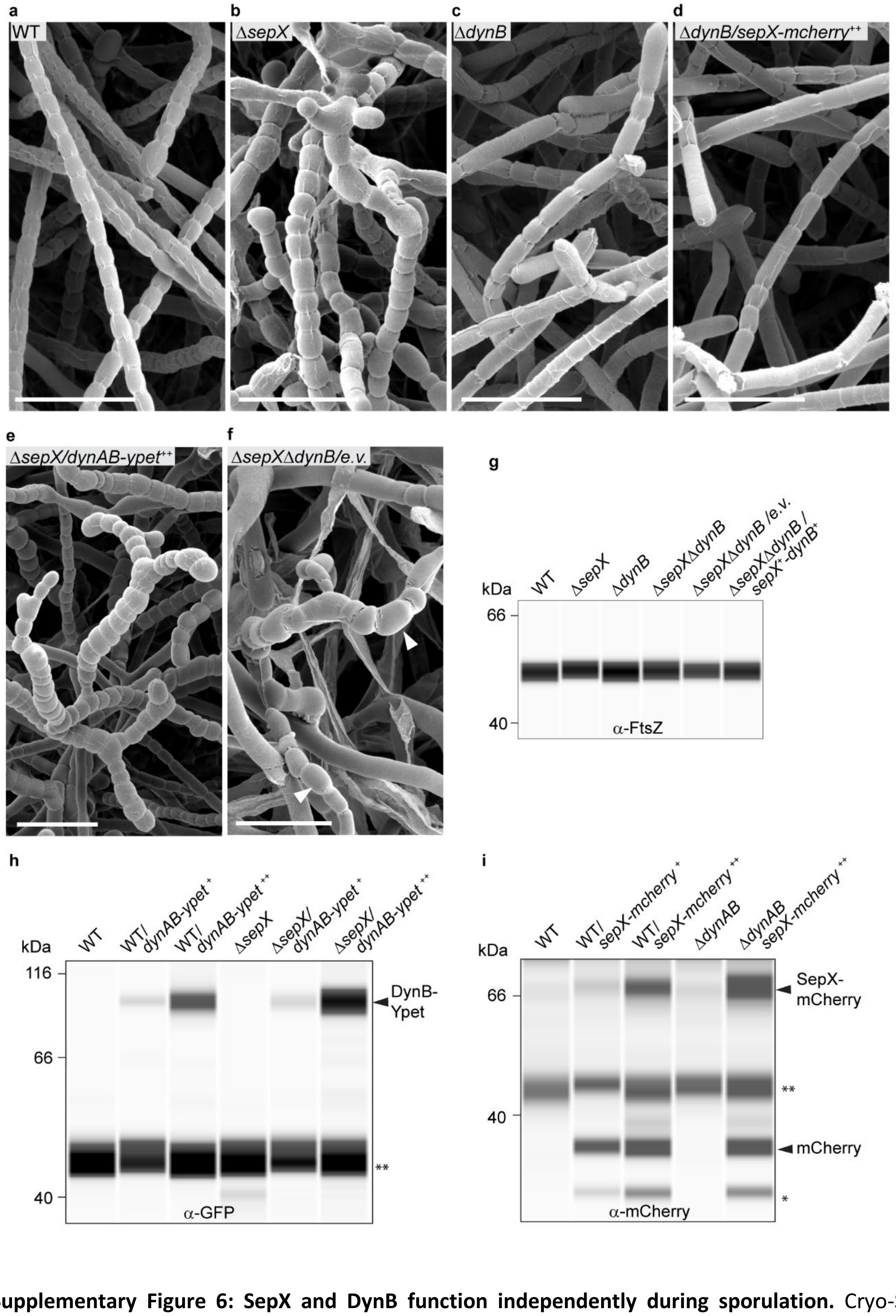

**Supplementary Figure 6: SepX and DynB function independently during sporulation.** Cryo-SEM images showing the terminal sporulation phenotype of **(a)** the wildtype (WT), **(b)** the Δ*sepX* mutant (SV55), **(c)** the Δ*dynB* mutant (SS2), **(d)** the Δ*dynB* mutant constitutively expressing *sepX-mcherry* (++, MB1092), **(e)** the Δ*sepX* mutant constitutively *dynAB-ypet* (++, MB1175) and **(f)** the Δ*sepX*Δ*dynB* double mutant carrying an empty vector (e.v., MB1099). White arrow heads in (f) point at the rare and aberrant spore chains. Scale bars: 5 μm. **(g)-(i)** Virtual Western blots showing the abundance of **(g)** FtsZ in Δ*sepX* (SV55), Δ*dynB* (SS2), Δ*sepX*Δ*dynB* (SV57) and the complemented mutant strain expressing both *sepX* and *dynB* (MB1103) or carrying the empty plasmid (MB1099); **(h)** DynB-YPet produced either from the native (+) or a constitutive (++) promoter in the wildtype (WT), SS92, MB1180, SV55, SS155 and MB1175 strains; **(i)** SepX-mCherry produced either from the native (+) or a constitutive (++) promoter in the WT, MB170, MB1124, LUV001 and MB1092 strains. Protein levels were verified using α-FtsZ, α-GFP and α-mCherry antibodies respectively. Asterisks denote unspecific cross-reactions of the antibody (**) or degradation products (*). Data shown in (a)-(f) are representative of at least seven images per strain. Western blots shown in (g)-(i) were performed in duplicate.

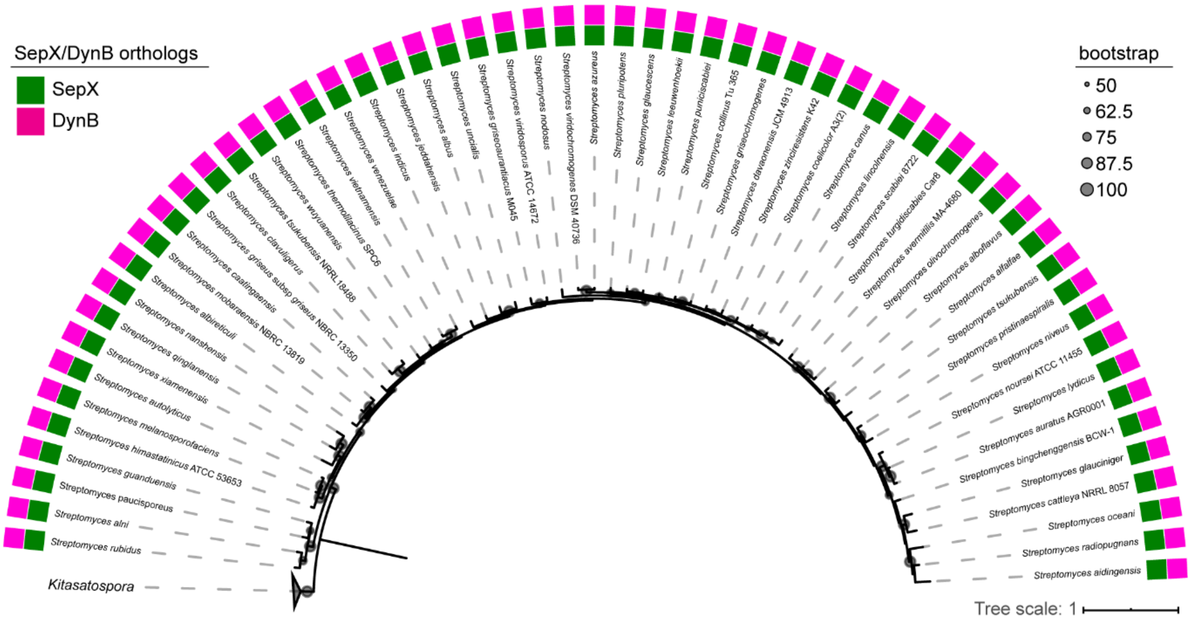

**Supplementary Figure 7: SepX and DynB are highly co-conserved in *Streptomyces*.** Distribution of SepX and DynB homologs in the genus *Streptomyces*. Maximum likelihood phylogeny of 58 representative *Streptomyces* genomes, based on concatenated alignment of 37 housekeeping genes. Tree is derived from the analysis presented in panel (b) with the *Streptomyces* clade un-collapsed. Bootstrap values >50% are indicated by grey circles at each node. Green and magenta boxes are used to represent the presence of SepX/DynB orthologs within each genome. Units of the tree are substitutions per site.

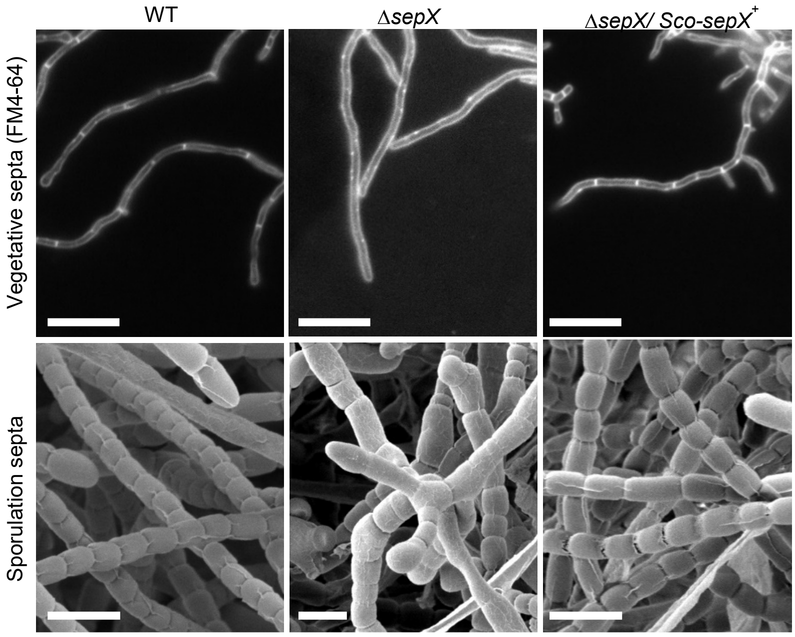

**Supplementary Figure 8.** Cross-complementation experiment. Expression of *sepX* from *S. coelicolor in trans* in the *S. venezuelae* Δ*sepX* mutant (MB1104) can restore vegetative septa formation and wildtype-like sporulation in the Δ*sepX* mutant (SV55). Vegetative septa were visualized using the fluorescent membrane dye FM4-64 (scale bars 10 μm). Spore chains were images by cryo-SEM (scale bars: 2 μm). Representative of at least five images are shown.

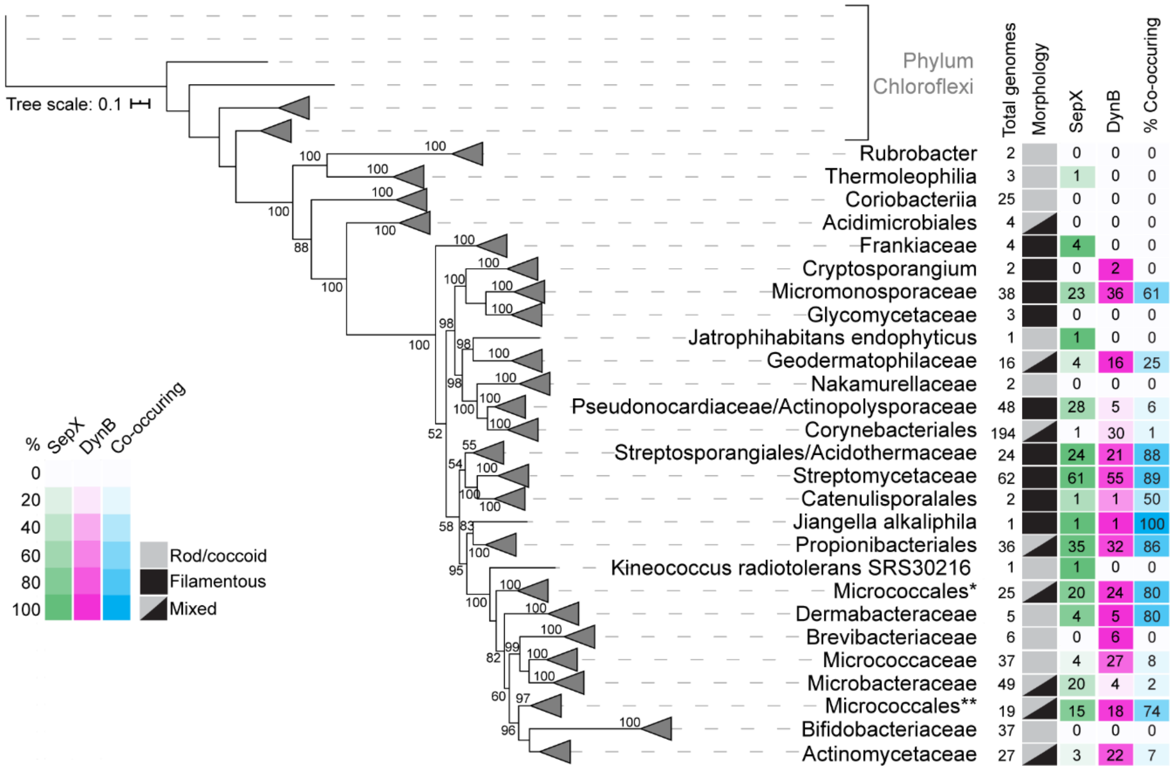

**Supplementary Figure 9: Distribution of SepX and DynB homologs in the Phylum Actinobacteria.** Data is presented as a maximum likelihood phylogenetic tree based on 673 representative actinobacterial genomes. Ten genomes from Phylum Chloroflexi are included as outgroups. Bootstrap values ≥50% are labeled at their respective nodes (values from 50-74% are indicated with †, 75-89% with ‡, and 90-100 with •). Triangles are used to represent where clades have been collapsed. For each clade, the total number of genomes, the number of genomes possessing a SepX and/or DynB ortholog, and the percentage of genomes where SepX/DynB co-occur are provided. Heatmaps are used to represent the percentage of total genomes within each clade possessing a SepX/DynB ortholog. Morphology designations for each clade are based on literature reports. *Includes Micrococcales families Dermatophilaceae, Intrasporangiaceae, Dermacoccaceae. **Includes Micrococcales families Promicromonosporaceae, Cellulomonadaceae, Sanguibacteraceae, Jonesiaceae, Ruaniaceae, and Beutenbergiaceae. Units of the tree are substitutions per site.

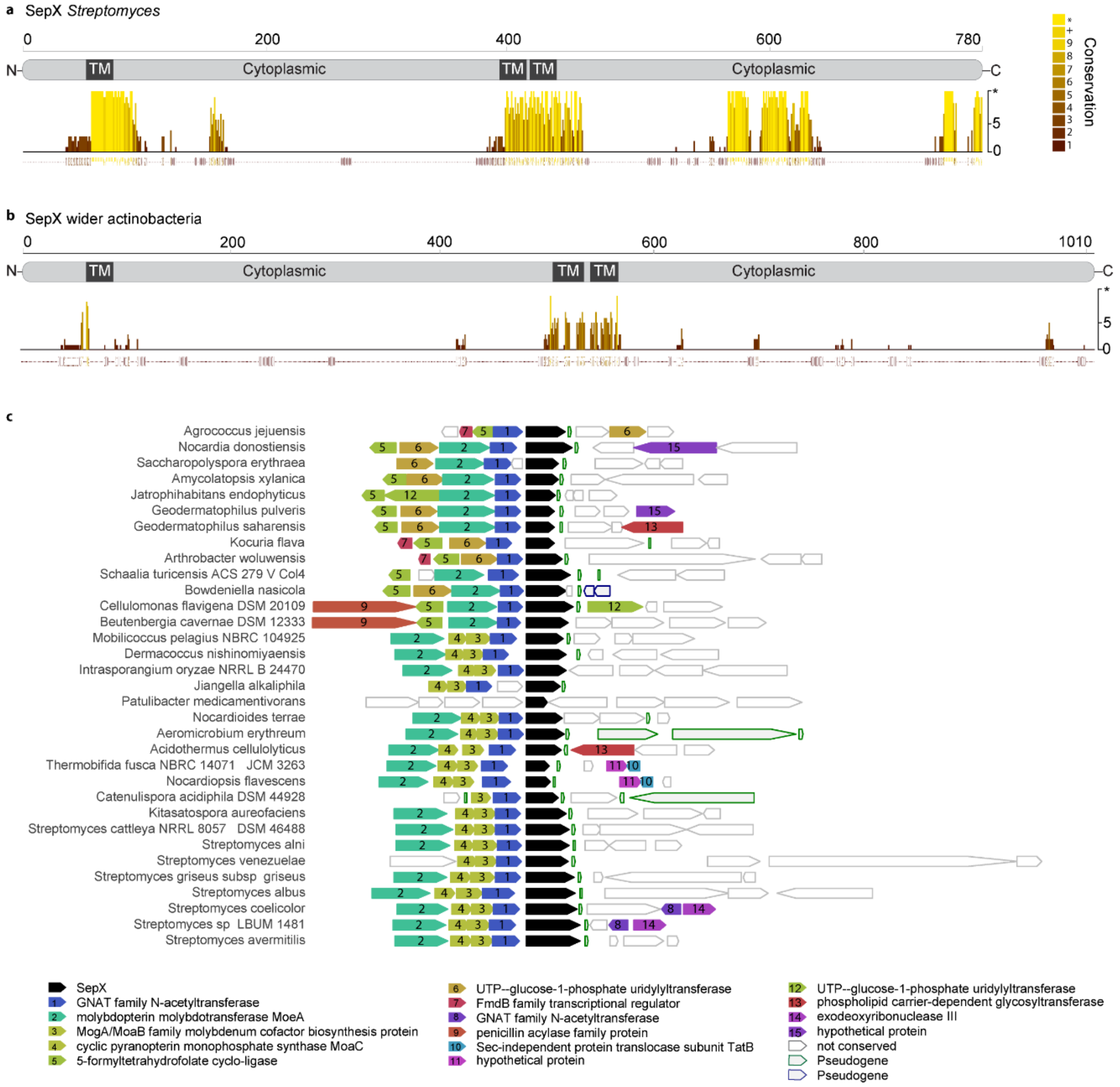

**Supplementary Figure 10: Sequence and gene neighborhood conservation of SepX homologs. (a-b)** Sequence conservation of SepX homologs from (a) *Streptomyces* (n=58) and (b) the wider actinobacteria (n=251). Conservation was calculated based on the Analysis of Multiply Aligned Sequences (AMAS) method implemented in Jalview[39]. **(c)** Gene neighborhood of *sepX* (shown in black) from 32 well-distributed representatives actinobacterial genomes that were selected based on phylogeny.

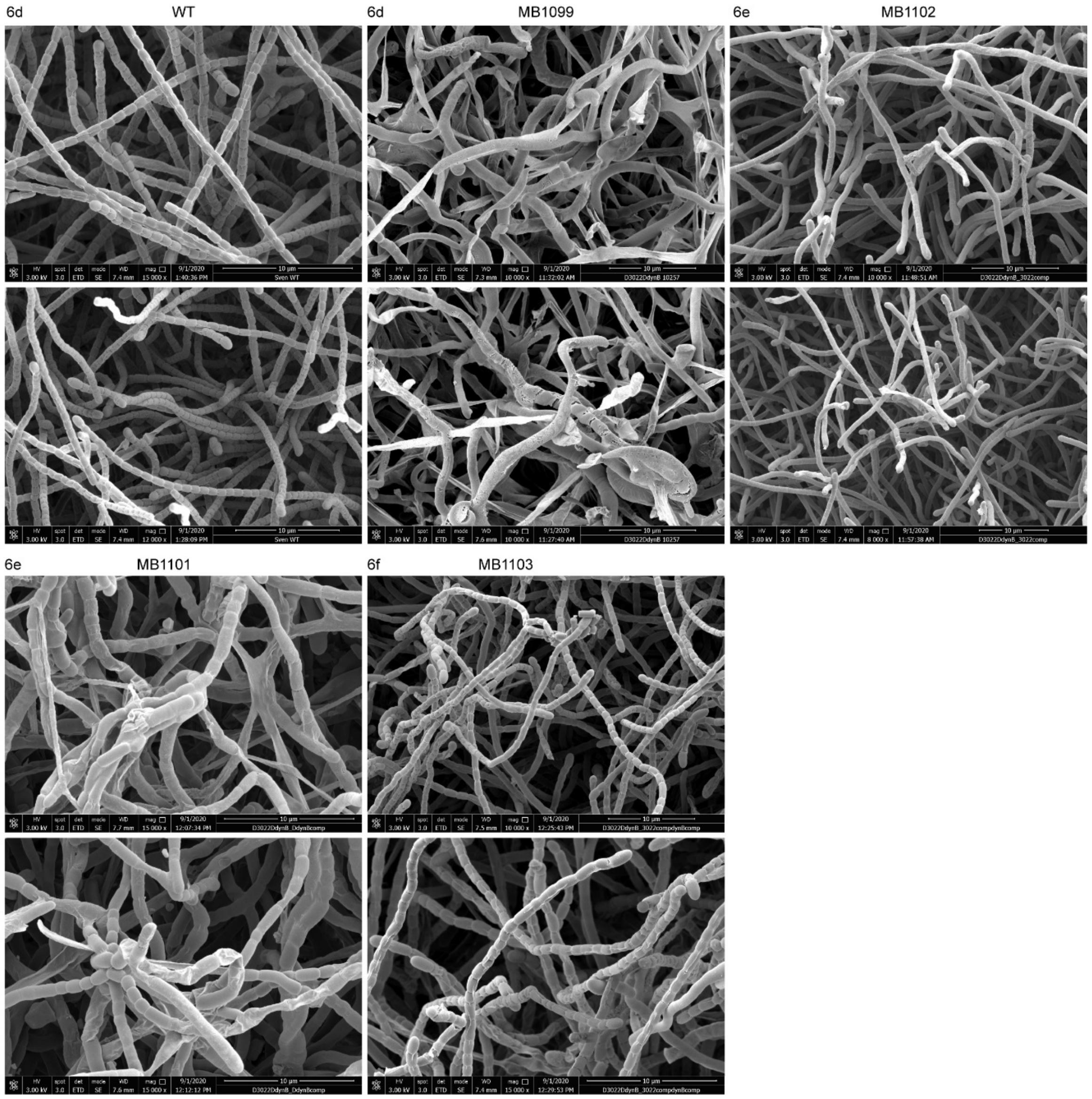

**Supplementary Figure 11: Full uncropped SEM images shown in Figure 6.** Numbers correspond to figure panels shown in Figure 6, additional representative SEMs for each strain are shown below.

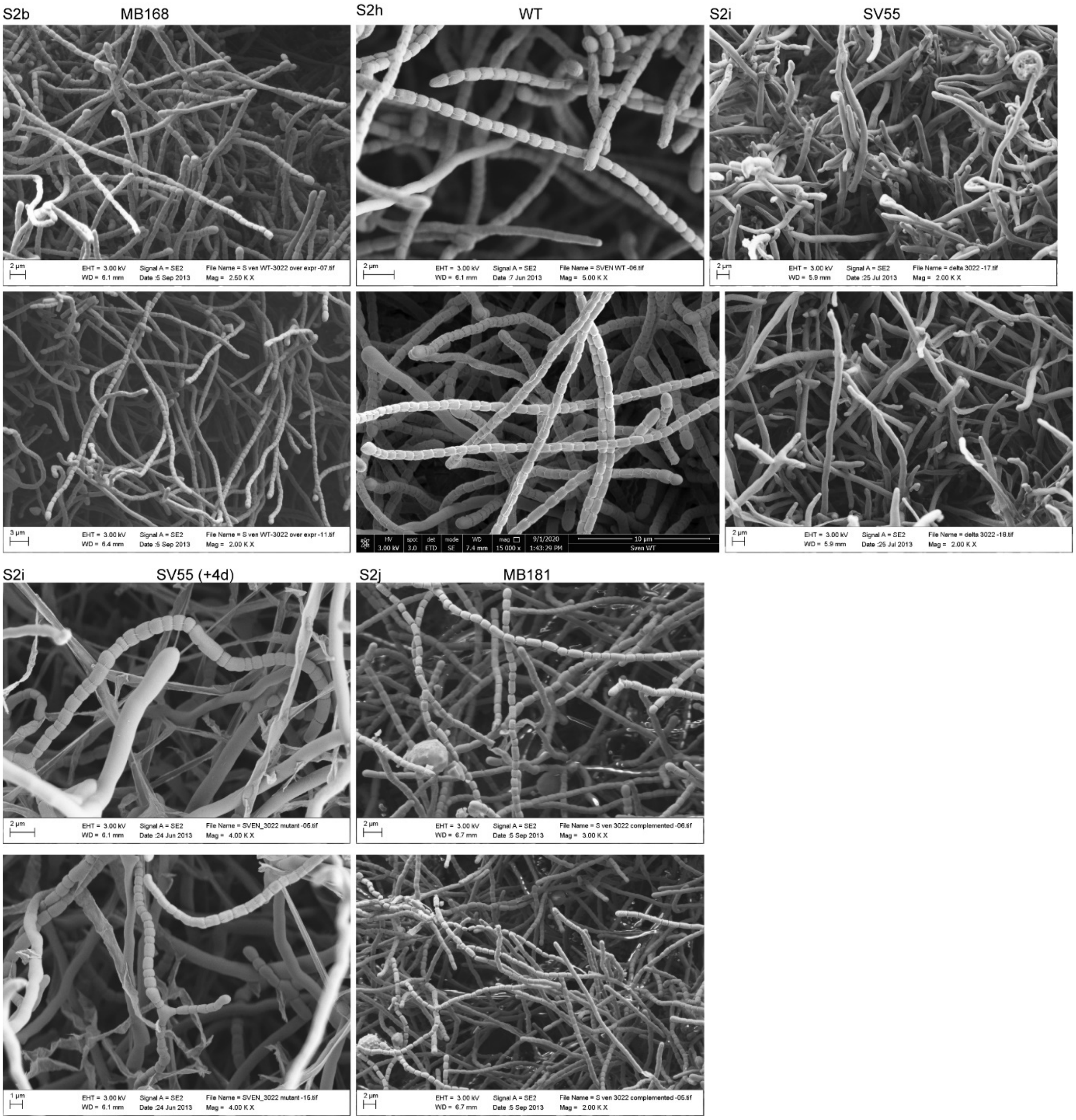

**Supplementary Figure 12: Full uncropped SEM image and additional SEMs corresponding to Supplementary Figure 2.** Note that the first SEM images corresponding to Supplementary Figure 2b was reflected vertically and the first image showing SEM S2j was rotated by 90$^{o}$.

S3e MB1120 MB171

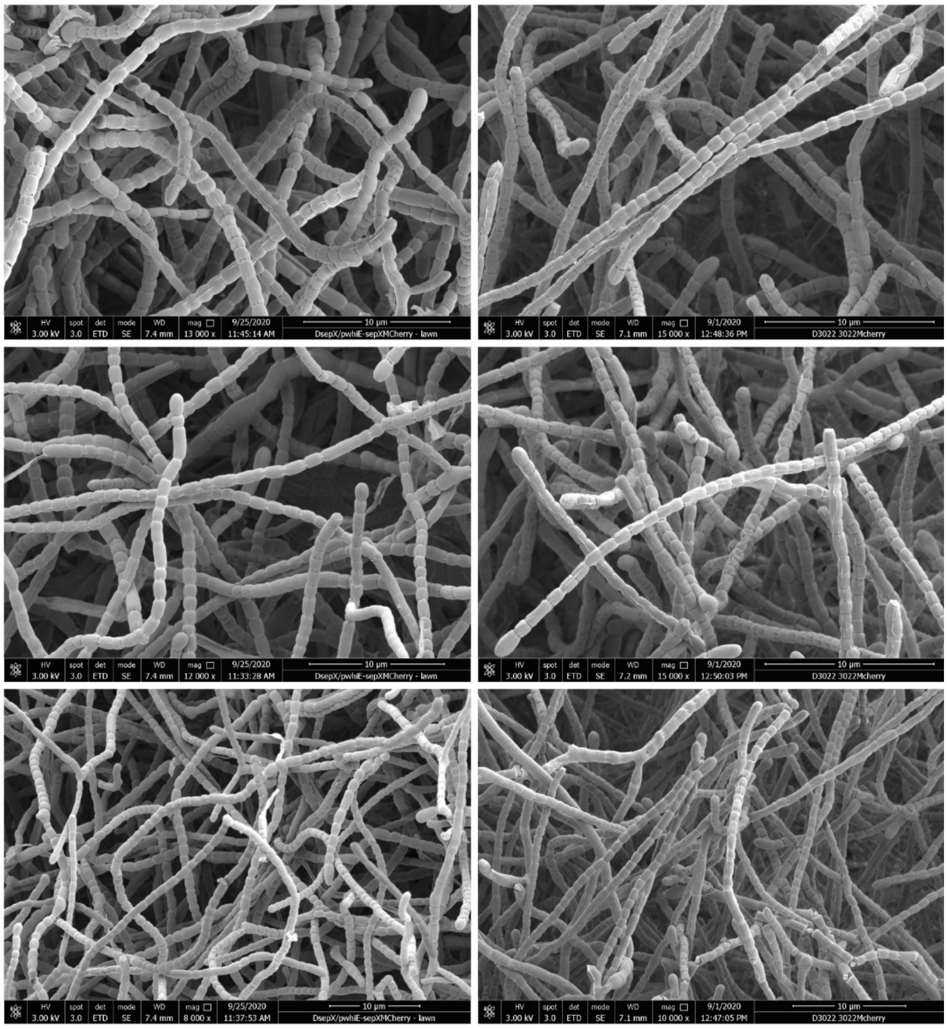

**Supplementary Figure 13: Full uncropped SEM images and additional SEMs corresponding to Supplementary Figure 3.** Note that the first SEM images corresponding to Supplementary Figure 3e was reflected vertically.

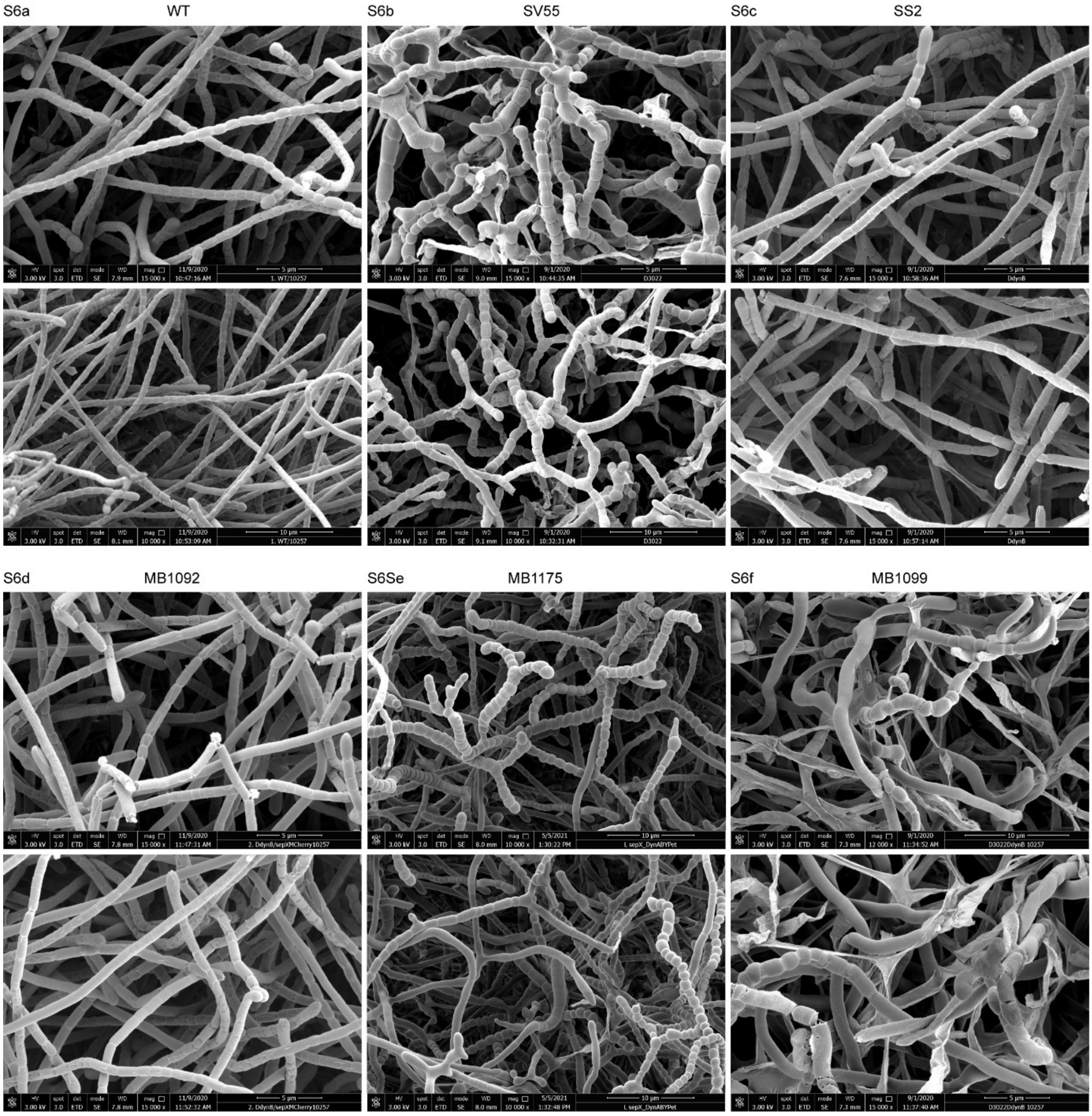

**Supplementary Figure 14: Full uncropped SEM images corresponding to Supplementary Figure 6.** For each strain, an additional representative SEM image is shown below.

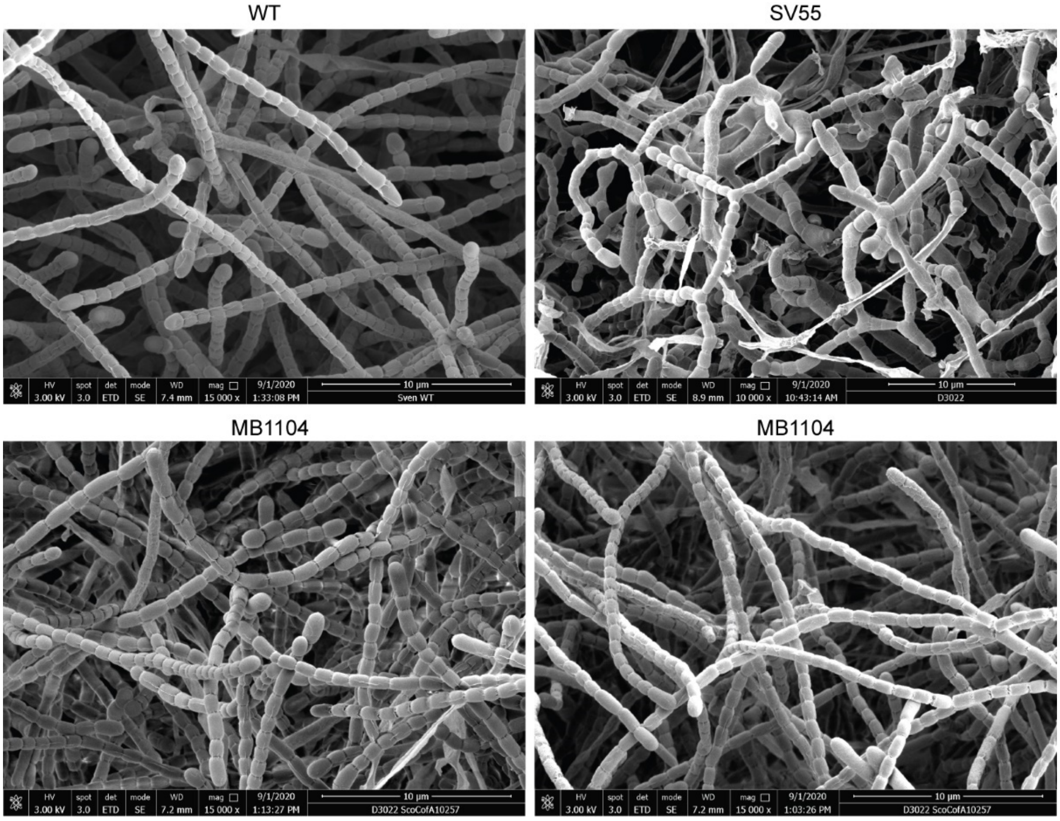

**Supplementary Figure 15: Full uncropped SEM images corresponding to Supplementary Figure 8.** For MB1104, an additional representative SEM image is shown.

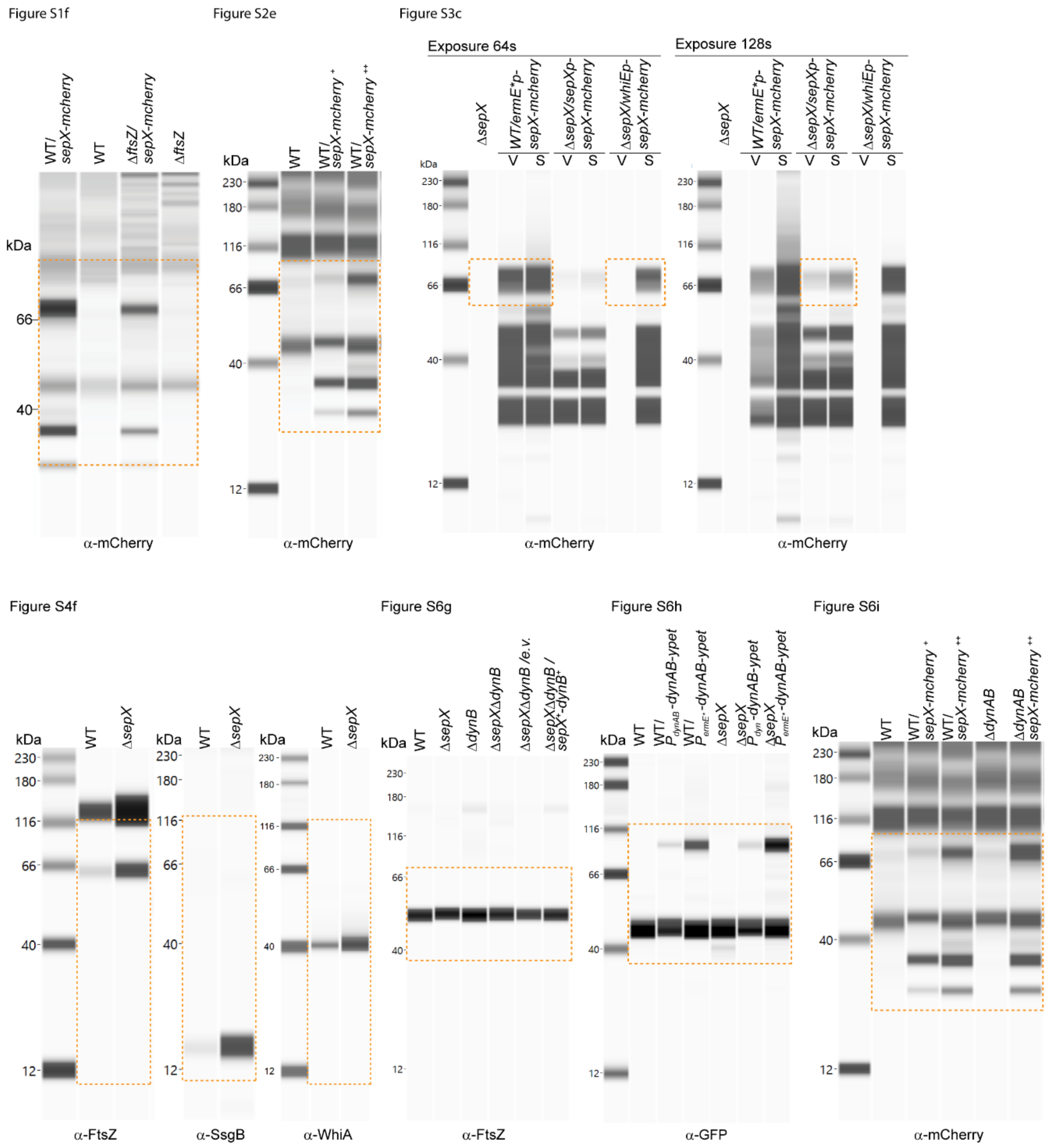

**Supplementary Figure 16: Full uncropped virtual Western blots of all the analysis shown in this work.** The orange boxes correspond to the crops used in the figures.

**Supplementary Table 1:** Strains used in this work.

| Strain | Details | Source | Reference |
|---|---|---|---|
| *S. venezuelae* | | | |
| NRRL B-65442 | Wild Type (WT) | Laboratory Strain | [2] |
| SV7 | Δ*whiA*::*apr* | | [3] |
| SV11 | Δ*whiB*::*apr* | | [4] |
| SV55 | Δ*sepX*::*apr* | Chromosomal *sepX* locus was replaced by *apra-oriT* by "redirect" PCR | This work |
| SV57 | Δ*sepX*::*hyg ΔdynB::apr* | Chromosomal *sepX* locus was replaced by *hyg-oriT* by "Redirect" in SS2 background | This work |
| DU669 | Δ*ftsZ::apr* | | [5] |
| LUV001 | Δ*dynAB::apr* | | [6] |
| LUV015 | WT *attB*$_{\Phi C31}$::*P*$_{ftsZ}$-*ftsZ-ypet* | | [6] |
| SS2 | Δ*dynB::apr* | | [6] |
| SS4 | WT *attB*$_{\Phi BT1}$::pIJ10257 | pIJ10257 integrated into the ΦBT1-attachment site of *S. venezuelae* | [6] |
| SS10 | Δ*dynAB::apr attB*$_{\Phi BT1}$::pIJ10257 | pIJ10257 integrated into the ΦBT1-attachment site of LUV001 | [6] |
| SS12 | WT a*ttB*$_{\Phi BT1}$::*P*$_{ftsZ}$-*ftsZ-ypet* | pSS5 integrated into the ΦBT1-attachment site of *S. venezuelae* | [6] |
| SS92 | WT *attB*$_{\phi BT1}$::*P*$_{dyn}$-*dynA-dynB-ypet* | pSS89 integrated into the ΦBT1-attachment site of *S. venezuelae* | This work |
| SS155 | Δ*sepX::apr attB*$_{\phi BT1}$::*P*$_{dyn}$-*dynA-dynB-ypet* | pSS89 integrated into the ΦBT1-attachment site of SV55 | This work |
| SS414 | WT *attB*$_{\Phi BT1}$::*P*$_{ermE*}$-*sepF-3xFLAG* | pSS559 integrated into the ΦBT1-attachment site of *S. venezuelae* | This work |
| MB127 | WT *attB*$_{\Phi BT1}$::*P*$_{ermE*}$-*ftsZ* | pSS3 integrated into the ΦBT1-attachment site of *S. venezuelae* | This work |
| MB168 | WT *attB*$_{\Phi BT1}$::*ermE*-sepX* | pMB156 integrated into the ΦBT1-attachment site of *S. venezuelae* | This work |
| MB170 | *WT attB*$_{\Phi BT1}$:: P$_{sepX}$-*sepX-mcherry* | pMB192 integrated into the ΦBT1-attachment site of *S. venezuelae* | This work |
| MB171 | Δ*sepX::apr attB*$_{\Phi BT1}$:: *P*$_{sepX}$-*sepX-mcherry* | pMB192 integrated into the ΦBT1-attachment site of SV55 | This work |
| MB180 | Δ*sepX::apr attB*$_{\Phi BT1}$::*P*$_{ftsZ}$-*ftsZ-ypet* | pSS5 integrated into the ΦBT1-attachment site of SV55 | This work |
| MB181 | Δ*sepX::apr attB*$_{\Phi BT1}$::*P*$_{sepX}$-*sepX* | pMB182 integrated into the ΦBT1-attachment site of SV55 | This work |
| MB182 | Δ*sepX::apr attB*$_{\Phi BT1}$::*P*$_{ermE*}$-*sepX* | pMB156 integrated into the ΦBT1-attachment site of SV55 | This work |
| MB192 | Δ*sepX::apr attB*$_{\Phi BT1}$::pMS82 | pMS82 integrated into the ΦBT1-attachment site of SV55 | This work |
| MB256 | WT *attB*$_{\Phi BT1}$::*P*$_{sepX}$-*sepX-mcherry attB*$_{\Phi C31}$::*P*$_{ftsZ}$-*ftsZ-ypet* | pMB192 integrated into the ΦBT1-attachment site and pKF351 integrated into the ΦC31-attachment site of *S. venezuelae* | This work |

| Strain | Details | Source | Reference |
|---|---|---|---|
| MB745 | Δ*sepX::apr* $attB_{\Phi BT1}$::$P_{sepX}$-*sepX* | pMB556 integrated into the ΦBT1-attachment site of SV55 | This work |
| MB839 | Δ*ftsZ::apr* $attB_{\Phi BT1}$::pIJ10257 | pIJ10257 integrated into the ΦBT1-attachment site of DU669 | This work |
| MB862 | Δ*sepX::apr* $attB_{\Phi BT1}$::pIJ10257 | pIJ10257 integrated into the ΦBT1-attachment site of SV55 | This work |
| MB942 | Δ*sepX::apr* $attB_{\Phi BT1}$::$P_{sepX}$-*sepX-3xFLAG*/$P_{dynAB}$-*dynA-dynB-ypet* | pMB703 integrated into the ΦBT1-attachment site of SV55 | This work |
| MB1104 | Δ*sepX::apr* $attB_{\Phi BT1}$::$P_{ermE*}$-*Sco-sepX* | pMB741 integrated into the ΦBT1-attachment site of SV55 | This work |
| MB1082 | Δ*ftsZ::apr* $attB_{\Phi BT1}$::$P_{ermE*}$-*sepX-mcherry* | pMB748 integrated into the ΦBT1-attachment site of DU669 | This work |
| MB1092 | Δ*dynAB::apr* $attB_{\Phi BT1}$:: $P_{ermE*}$-*sepX-mcherry* | pMB748 integrated into the ΦBT1-attachment site of LUV001 | This work |
| MB1099 | Δ*sepX::hyg* Δ*dynB::apr* $attB_{\Phi BT1}$::*pIJ10257* | pMB743 integrated into the ΦBT1-attachment site of SV57 | This work |
| MB1101 | Δ*sepX::hyg* Δ*dynB::apr* $attB_{\Phi BT1}$:: *ermE*-dynB* | pMB745 integrated into the ΦBT1-attachment site of SV57 | This work |
| MB1102 | Δ*sepX::hyg* Δ*dynB::apr attBΦBT1:: ermE*-sepX* | pMB744 integrated into the ΦBT1-attachment site of SV57 | This work |
| MB1103 | Δ*sepX::hyg* Δ*dynB::apr* $attB_{\Phi BT1}$:: $P_{ermE*}$-*dynB*; $P_{sepX}$-*sepX* | pMB746 integrated into the ΦBT1-attachment site of SV57 | This work |
| MB1111 | Δ*sepX::hyg* Δ*dynB::apr* $attB_{\Phi BT1}$::*pKF351* | pKF351 integrated into the ΦBT1-attachment site of SV57 | This work |
| MB1120 | Δ*sepX::apr* $attB_{\Phi BT1}$:: $P_{whiE}$-*sepX-mcherry* | pSS601 integrated into the ΦBT1-attachment site of SV55 | This work |
| MB1124 | WT $attB_{\Phi BT1}$::$P_{ermE*}$-*sepX-mcherry* | pMB748 integrated into the ΦBT1-attachment site of *S. venezuelae* | This work |
| MB1175 | Δ*sepX::apr* $attB_{\phi BT1}$::$P_{ermE*}$-*dynA-dynB-ypet* | pSS134 integrated into the ΦBT1-attachment site of SV55 | This work |
| MB1180 | WT $attB_{\phi BT1}$:: $P_{ermE*}$-*dynA-dynB-ypet* | pSS134 integrated into the ΦBT1-attachment site of *S. venezuelae* | This work |
| MB1270 | Δ*dynAB*::*apr* $attB_{\Phi BT1}$::$P_{sepX}$-*sepX-mcherry* | pMB192 integrated into the ΦBT1-attachment site of LUV001 | This work |
| *E.coli* | | | |
| ET12567(pUZ8002) | *F⁻ dam13::Tn9 dcm6 hsdM hsdR recF143::Tn10 galK2 galT22 ara-14 lacY1 xyl-5leuB6 thi-1 tonA31 rpsL hisG4 tsx-78 mtl-1 glnV44* | ET12567 containing helper plasmid pUZ8002 | [7] |
| BW25113 | Δ(*araD-araB*)*567* Δ *lacZ4787*(::*rrnB-4*) *lacIp-4000*($lacI^{Q}$), l-*rpoS369*(*Am*) *rph-1* Δ(*rhaD-rhaB*)*568 hsdR514* | BW25113 containing λ RED recombination plasmid | [8] |
| BTH101 | *F⁻ cya-99 araD139 galE15 galK16 rpsL1* ($Str^{R}$) *hsdR2 mcrA1 mcrB1* | Bacterial two-hybrid host strain | [9] |

**Supplementary Table 2:** Plasmids used in this work.

| Plasmid | Details | Construction | Reference |
|---|---|---|---|
| pIJ773 | pBluescript KS (+) containing the apramycin resistance gene *apr* and *oriT* of plasmid RP4, flanked by FRT sites ($Apr^R$). Used as template for the amplification of the *apr oriT* cassette for 'Redirect' PCR-targeting | | [10] |
| pIJ10700 | pBluescript KS (+) containing the hygromycin resistance gene *hyg* and *oriT* of plasmid RP4, flanked by FRT sites ($Apr^R$). Used as template for the amplification of the *hyg oriT* cassette for 'Redirect' PCR-targeting | | [9] |
| pIJ790 | Modified λ RED recombination plasmid *[oriR101] [repA101*(ts)*] araBp-gam-bet-exo* ($Cam^R$) | | [9] |
| pIJ10754 | pUC19-derivative used to generate 3xFLAG gene fusions ($Amp^R$) | | [6] |
| pIJ10750 | pMS82 with an extended Multiple Cloning Site, ($Hyg^R$) | | [5] |
| pIJ10770 | Modified pIJ10750 lacking an intrinsic apramycin promoter upstream of the extended multiple cloning site ($Hyg^R$) | | [5] |
| pIJ10257 | Plasmid cloning vector for the conjugal transfer of DNA (under control of the *ermE** constitutive promoter) from *E. coli* to *Streptomyces spp.* Integrates specifically at the ΦBT1 attachment site ($Hyg^R$) | | [11] |
| pIJ6772 | pKT25 carrying *rsbN* ($Kan^R$) | | [12] |
| pIJ6769 | pUT18C carrying *bldN* ($Amp^R$) | | [11] |
| pMS82 | Plasmid cloning vector for the conjugal transfer of DNA from *E. coli* to *Streptomyces* spp. Integrates site specifically at the ΦBT1 attachment site ($Hyg^R$) | | [13] |
| pTB145 | Plasmid for overexpression of His6-Upl1 protease ($Amp^R$) | | [14] |

| Plasmid | Details | Construction | Reference |
|---|---|---|---|
| pTB146 | Plasmid for overexpression of proteins with N-terminal $His_6$-SUMO fusion ($Amp^R$) | | [14] |
| pKF280 | Vector for the construction of gene fusions to *ypet*, based on pIJ6902 but with the *tipA* promoter removed. ($Apr^R$) | | [15] |
| pKF351 | Derivative of pKF280 encoding the FtsZ-YPet fusion ($Apr^R$) | | [6] |
| pKT25 | Two-hybrid plasmid, N-terminal *cyaAT25* fusion ($Kan^R$) | | [9] |
| pKNT25 | Two-hybrid plasmid, C-terminal *cyaAT25* fusion ($Kan^R$) | | [8] |
| pUT18 | Two-hybrid plasmid, C-terminal *cyaAT18* fusion ($Amp^R$) | | [8] |
| pUT18C | Two-hybrid plasmid, N-terminal *cyaAT18* fusion ($Amp^R$) | | [8] |
| pSS3 | pIJ10257 carrying *ftsZ* ($Hyg^R$) | | [6] |
| pSS5 | pIJ10750 carrying *pftsZ-ftsZ-ypet (*$Hyg^R$) | | [6] |
| pSS23 | pKT25 carrying *dynA* ($Kan^R$) | *dynA* (*vnz_12110*) was PCR amplified using primers ss65/66 and inserted into pKT25 via restriction cloning using EcoRI/BamHI. | This work |
| pSS64 | pIJ10257 carrying *dynB* ($Hyg^R$) | *dynB (vnz_12105)* was PCR amplified using primers ss21/97 and inserted into pIJ10257 via restriction cloning using NdeI/HindIII. | This work |
| pSS89 | pKF227 carrying $P_{dynAB}$-*dynA-dynB-ypet* ($Hyg^R$) | | [6] |
| pSS105 | pKT25 carrying *dynB* ($Kan^R$) | | [6] |
| pSS115 | pUT18C carrying *dynB* ($Amp^R$) | | [6] |
| pSS125 | pKT25 carrying *dynB*Δ468-517 ($Kan^R$) | *dynB*[illegible]*TM* (ΔAA468-517) was generated by amplifying the two *dynB* fragments using primers ss69/190 and ss70/189, followed by restriction digestion with EcoRI/Asp718 and Asp718/BamHI. Both fragments were ligated between the EcoRI-BamHI site of pKT25. | This work |

| Plasmid | Details | Construction | Reference |
|---|---|---|---|
| pSS133 | pIJ10257 carrying *dynB-3xFLAG* ($Hyg^R$) | *dynB* was amplified using primer ss105/106 and first cloned into pIJ10754 between the KpnI-XhoI site. The *dynB-3xFLAG* fragment was then amplified with primer ss209/21 and subcloned into pIJ10257 between the NdeI-HindIII site. | This work |
| pSS134 | pIJ10257 carrying *dynA-dynB-ypet* ($Hyg^R$) | | [6] |
| pSS172 | pIJ10750 carrying *mcherry* ($Hyg^R$) | | [6] |
| pSS196 | | *sepF (vnz_08505)* was PCR amplified using primer ss336/337 and inserted into pSS172 via restriction cloning using HindIII/AvrII. | This work |
| pSS220 | pKT25 carrying *sepF2* ($Kan^R$) | | [6] |
| pSS221 | pKT25 carrying *sepF3* ($Kan^R$) | | [6] |
| pSS222 | pKT25 carrying *sepF* ($Kan^R$) | | [6] |
| pSS440 | pTB146 carrying ssgB ($Carb^R$) | *ssgB (vnz_05545)* was PCR amplified with primer ss889/ss890, followed by gibson assembly into pTB146 cut with BpsQI | This work |
| pSS559 | pIJ10257 carrying *sepF-3xFLAG* ($Hyg^R$) | *sepF (vnz_08505)* was PCR amplified from pSS196 with primer ss418/ss5, followed by restriction digestion with NdeI/XhoI; *3xFLAG* fragment was isolated from pSS133 cut with NcoI/XhoI; pIJ10257 was cut with NcoI/NdeI; all three fragments were assembled by ligation. | This work |
| pSS601 | pMB748 modified so that *sepX-mcherry* under the control of $P_{whiE}$ ($Hyg^R$) | *whiE* promoter region was PCR amplified with primer ss1372/ss1373 and inserted via Gibson Assembly into pMB748 cut with Bsu36I and NdeI to remove *ermE** promoter. | This work |
| pMB156 | pIJ10257 carrying *sepX* ($Hyg^R$) | *sepX* was PCR amplified using primers mb331/mb332 and inserted into pIJ10257 via restriction cloning using NdeI/HindIII. | This work |

| Plasmid | Details | Construction | Reference |
|---|---|---|---|
| pMB174 | pKT25 carrying *ssgB* ($Kan^R$) | *ssgB* was PCR amplified using primers mb364/mb365 and inserted into pKT25 via restriction cloning using BamHI/KpnI. | This work |
| pMB177 | pKNT25 carrying *sepX* ($Kan^R$) | *sepX* was PCR amplified using primers mb366/mb367 and inserted into pKNT25 via restriction cloning using XbaI/BamHI. | This work |
| pMB180 | pUT18 carrying *sepX* ($Amp^R$) | *sepX* was PCR amplified using primers mb366/mb367 and inserted into pUT18 via restriction cloning using XbaI/BamHI. | This work |
| pMB182 | pIJ10750 carrying $P_{sepX}$-*sepX* ($Hyg^R$) | $P_{sepX}$-*sepX* was PCR amplified using primers mb210/mb340 and inserted into pIJ10750 via restriction cloning using HindIII/AvrII. | This work |
| pMB192 | pIJ10750 carrying $P_{sepX}$-*sepX-mcherry* ($Hyg^R$) | Two-step amplification using mb210/mb382 and mb384/mb385 in the first step. Templates from first step amplified using mb210 and mb385 to generate $P_{sepX}$-*sepX -mcherry* fragment which was inserted into pIJ10750 via restriction cloning using HindIII/AvrII. | This work |
| pMB555 | pIJ10770 carrying $P_{sepX}$-*sepX-FLAG* ($Hyg^R$) | Two-step amplification using mb373/mb377 and mb378/mb374 in the first step. Templates from first step amplified using mb373 and mb374 to generate $P_{sepX}$-*sepX-FLAG* fragment which was inserted into pIJ10770 via restriction cloning using HindIII/AvrII. | This work |
| pMB556 | pIJ10770 carrying $P_{sepX}$-*sepX* ($Hyg^R$) | $P_{sepX}$-*sepX* was PCR amplified using mb1433/mb1434 and pMB182 as a template and cloned into EcoRV-cut pIJ10770 using Gibson Assembly. | This work |
| pMB703 | pSS89 carrying $P_{sepX}$-*sepX-3xFLAG* ($Hyg^R$) | $P_{sepX}$-*sepX-3xFLAG* was PCR amplified using mb1150/mb1151 and pMB555 as a template and cloned into EcoRV-cut pSS89 using Gibson Assembly. | This work |
| pMB741 | pIJ10257 carrying *S. coelicolor sepX* | *Sco3177* was PCR amplified using mb1437/mb1438 and cosmid stE87 as a template and inserted into pIJ10257 via restriction cloning using NdeI/HindIII. | This work |

| Plasmid | Details | Construction | Reference |
|---|---|---|---|
| pMB744 | pMB156 carrying additional *thio*$^R$ | Thiostrepton resistance cassette was amplified from pKF351 with primer mb1451/mb1452 followed by Gibson Assembly into EcoRV-cut pMB156. | This work |
| pMB745 | pSS64 carrying additional *thio*$^R$ | Thiostrepton resistance cassette was amplified from pKF351 with primer mb1451/mb1452 followed by Gibson Assembly into EcoRV-cut pSS64. | This work |
| pMB746 | pMB745 carrying $P_{sepX}$-*sepX* | $P_{sepX}$-*sepX* was amplified using primers mb1453/mb1454 and pMB192 as template and assembled into SpeI-cut pMB745. | This work |
| pMB748 | pIJ10257 carrying *sepX-mcherry* | *sepX-mcherry* was PCR amplified using primers mb331/mb1448 and pMB192 as template and inserted into pIJ10257 via restriction cloning using NdeI/HindIII. | This work |

**Supplementary Table 3:** Primer used in this work.

| Primer | Sequence |
|---|---|
| mb210 | GGCGAAGCTTCCCGCTACCTGCACATCG |
| mb266 | TGCCGTCCCGCGAACCCCTCTACCGTGTGAGGCGTGAGCATTCCGGGGATCCGTCGACC |
| mb267 | CAGCCGGGGGCCGTCGCGGCGCCCCCGGCGGGGTCATTCTGTAGGCTGGAGCTGCTTC |
| mb331 | GGAATTCCATATGAGCAGCAGCGGCCTCA |
| mb332 | CCCAAGCTTTCATTCGTTGGCCGCCCG |
| mb340 | GCTGCCTAGGTCATTCGTTGGCCGCCCG |
| mb364 | CTGAGGATCCCATGAACACCACGGTCAGCTG |
| mb365 | CCGGTACCCGGCTGTCGGCCAGGATGTG |
| mb366 | GCTCTAGAGGTGAGCAGCAGCGGCCTC |
| mb367 | CGCGGATCCTCTTCGTTGGCCGCCCGGG |
| mb373 | GGCGAAGCTTCCCGCTACCTGCACATCGACG |
| mb374 | GCTGCCTAGGCCACAGGCCCTTGCGACGTG |
| mb377 | TCGATGTCGTGGTCCTTGTAGTCGCCGTCGTGGTCCTTGTAGTCTTCGTTGGCCGCCCGGGG |
| mb378 | CGACTACAAGGACCACGACATCGACTACAAGGACGATGACGACAAGTGACCCCGCCGGGGGCG |
| mb382 | CTTGGAGACTTCGTTGGCCGCCCGGGG |
| mb384 | GCCAACGAAGTCTCCAAGGGCGAGGAG |
| mb385 | GCTGCCTAGGTCACTTGTACAGCTCGTCCATG |
| mb1150 | GGAGCGCGGCCGCGCGCGATCCCGCTACCTGCACATCG |
| mb1151 | GACATGATTACGAATTCGATCCACAGGCCCTTGCGACG |
| mb1431 | GTGCCGGAGGGGCTGCTG |
| mb1432 | GCATCCCGACCGGCTCAGG |
| mb1433 | GCGGCCGCGCGCGATCCCGCTACCTGCACATCG |
| mb1434 | ACATGATTACGAATTCGATTCATTCGTTGGCCGCCCG |
| mb1437 | GGGAATTCCATATGAGCAGCAGCGGCCTCATCT |
| mb1438 | CCCAAGCTTCTACTCGTTGGCCGCGCGG |
| mb1448 | CCCAAGCTTTCACTTGTACAGCTCGTCCATGCCG |

| Primer | Sequence |
|---|---|
| mb1451 | ACCATAGCGGGCAGGGAGCGGATCCGCGGCCGCGCGCGATATCGCTCATGAGCGGAGAACGAGATGACGTTGG AGGGGCAAGGTCGCG |
| mb1452 | TTCACACAGGAAACAGCTATGACATGATTACGAATTCGATATCCTTATCGGTTGGCCGCGAGATTCCT |
| mb1453 | TCGGCCCCTTTTTTGGCCTTGAAATCGTTAGTTAGGCTAACCCGCTACCTGCACATCGACG |
| mb1454 | GCGGATCCGCTCCCTGCCCGCTATGGTGACGAAGGAACTATCATTCGTTGGCCGCCCGG |
| ss5 | TGGCCGTTCACGGAGCCCTC |
| ss21 | ATTAATTCATATGGTGGATGTGGACGTGGAAGCG |
| ss65 | TTAAGGATCCCATGGTGACCTTGGACGAACGGCC |
| ss66 | AATGAATTCTCACTTCTCCTTCTGCAGTACGGAGAGC |
| ss69 | AATTAAGGATCCCATGGTGGATGTGGACGTGGAAGCG |
| ss70 | AATTAATGAATTCCTACCTGCCCCGTACCCCCGTC |
| ss97 | TTAATTCCTAGGCTACCTGCCCCGTACCCCCGTCC |
| ss105 | ATCGGGGTACCGTGGTGGATGTGGACGTGGAAGCG |
| ss106 | ATCCGCTCGAGCCTGCCCCGTACCCCCGTCCGCG |
| ss189 | ATCGGGGTACCCCGGTCCTCGTGATGGTCGGCG |
| ss190 | ATCGGGGTACCGACCGTCAGCTCGTCGAGCGCCTC |
| ss209 | GAGCGGATAACAATTTCACACAGG |
| ss336 | ATTAAAGCTTAGCTCGGCTGCCTCCTGCAGGTC |
| ss337 | TTAATTCCTAGGGCTCTGGTTGAAGAATCCGCCCTCTG |
| ss418 | AATTAATTCATATGGCCGGCGCGATGCGCAAG |
| ss889 | TGAGGCTCACAGAGAACAGATTGGTGGTAGAAGAGCAATGAACACCACGGTCAGCTG |
| ss890 | TTGTCGACGGAGCTCTGCTCTTCTATCAGCTGTCGGCCAGGATGT |
| ss1372 | CCTCGCCTCTGACCCCTGACCCCGTCAGCCTCCCCGGCCGAGTCGAATCGG |
| ss1373 | TGACTGCGTAGATGAGGCCGCTGCTGCTCATATGGCACACCTCTTCTCTTCTCGGTGGGGGGTGGGG |

### Supplementary References

1. Al-Bassam, M. M., Bibb, M. J., Bush, M. J., Chandra, G. & Buttner, M. J. Response regulator heterodimer formation controls a key stage in *Streptomyces* development. *PLOS Genetics* **10**, e1004554 (2014).
2. Gomez-Escribano, J. P. *et al. Streptomyces venezuelae* NRRL B-65442: genome sequence of a model strain used to study morphological differentiation in filamentous actinobacteria. *J Ind Microbiol Biotechnol* kuab035 (2021) doi:10.1093/jimb/kuab035.
3. Bush, M. J., Bibb, M. J., Chandra, G., Findlay, K. C. & Buttner, M. J. Genes required for aerial growth, dell division, and chromosome segregation are targets of WhiA before sporulation in *Streptomyces venezuelae*. *mBio* **4**, e00684-13 (2013).
4. Bush, M. J., Chandra, G., Bibb, M. J., Findlay, K. C. & Buttner, M. J. Genome-wide chromatin immunoprecipitation sequencing analysis shows that WhiB Is a transcription factor that cocontrols its regulon with WhiA to initiate developmental dell division in *Streptomyces*. *mBio* **7**, e00523-16, /mbio/7/2/e00523-16.atom (2016).
5. Santos-Beneit, F., Roberts, D. M., Cantlay, S., McCormick, J. R. & Errington, J. A mechanism for FtsZ-independent proliferation in *Streptomyces*. *Nat Commun* **8**, 1378 (2017).
6. Schlimpert, S. *et al.* Two dynamin-like proteins stabilize FtsZ rings during *Streptomyces* sporulation. *PNAS* **114**, E6176–E6183 (2017).

7. Paget, M. S. B., Chamberlin, L., Atrih, A., Foster, S. J. & Buttner, M. J. Evidence that the extracytoplasmic function sigma factor σE is required for normal cell wall structure in *Streptomyces coelicolor* A3(2). *J. Bacteriol.* **181**, 8 (1999).
8. Datsenko, K. A. & Wanner, B. L. One-step inactivation of chromosomal genes in *Escherichia coli* K-12 using PCR products. *PNAS* **97**, 6640–6645 (2000).
9. Karimova, G., Pidoux, J., Ullmann, A. & Ladant, D. A bacterial two-hybrid system based on a reconstituted signal transduction pathway. *PNAS* **95**, 5752–5756 (1998).
10. Gust, B., Challis, G. L., Fowler, K., Kieser, T. & Chater, K. F. PCR-targeted *Streptomyces* gene replacement identifies a protein domain needed for biosynthesis of the sesquiterpene soil odor geosmin. *PNAS* **100**, 1541–1546 (2003).
11. Hong, H.-J., Hutchings, M. I., Hill, L. M. & Buttner, M. J. The role of the novel Fem protein VanK in vancomycin resistance in *Streptomyces coelicolor*. *J. Biol. Chem.* **280**, 13055–13061 (2005).
12. Bibb, M. J., Domonkos, Á., Chandra, G. & Buttner, M. J. Expression of the chaplin and rodlin hydrophobic sheath proteins in *Streptomyces venezuelae* is controlled by σBldN and a cognate anti-sigma factor, RsbN. *Molecular Microbiology* **84**, 1033–1049 (2012).
13. Gregory, M. A., Till, R. & Smith, M. C. M. Integration site for *Streptomyces* phage phiBT1 and development of site-specific integrating vectors. *J. Bacteriol.* **185**, 5320–5323 (2003).
14. Bendezú, F. O., Hale, C. A., Bernhardt, T. G. & de Boer, P. A. J. RodZ (YfgA) is required for proper assembly of the MreB actin cytoskeleton and cell shape in *E. coli*. *EMBO J.* **28**, 193–204 (2009).
15. Donczew, M. *et al.* ParA and ParB coordinate chromosome segregation with cell elongation and division during *Streptomyces* sporulation. *Open Biol* **6**, 150263 (2016).
